# Non-parallel morphological divergence following colonization of a new host plant

**DOI:** 10.1101/2021.03.30.436984

**Authors:** Kalle J. Nilsson, Jesús Ortega, Magne Friberg, Anna Runemark

## Abstract

Divergent ecological selection may diversify conspecific populations evolving into different niches, and may lead to speciation if gene flow is ceased, e.g, due to reinforcement leading to character displacement. Adaptation to the same niche is expected to be parallel. Whether selection against maladaptive hybridization in secondary sympatry also results in parallel divergence in traits that are not directly related to the ecological niches remains an empirical challenge, although such character displacement can be decisive for if ecotypes develop into new species. Here, we use a host shift in the phytophagous peacock fly *Tephritis conura*, with both host races represented in two geographically separate areas East and West of the Baltic Sea to investigate convergence in morphological adaptations. We asked i) if there are consistent morphological adaptations to a host plant shift and ii) if the response to secondary sympatry with the alternate host race is parallel across contact zones. We found surprisingly low and variable, albeit significant, divergence between host races. Only one trait, the length of the female ovipositor, which serves an important function in the interaction with the hosts, was consistently different. Instead, co-existence with the other host race significantly affected the degree of morphological divergence, but the divergence was largely driven by different traits in different contact zones. Thus, local stochastic fixation or reinforcement could generate trait divergence, and additional evidence is needed to conclude whether divergence is locally adaptive.

## Introduction

Adaptation to novel niches through ecological selection has been identified as a major driver of speciation (Schluter 2008; Schluter 2009; Nosil 2012). Classic adaptive radiations including the Darwin Finches (Weiner 1994; Loo et al 2019), the Hawaiian tarweeds and silverswords (Baldwin et al 2003) and the recurrent evolution of limnetic-benthic stickleback pairs (Schluter 1993; Bay et al 2017) are driven by adaptations to novel diets and habitats. Speciation driven by expansion of novel dietary niches has potential to be rapid, like in *Rhagoletis* flies, where populations of hawthorn-feeding flies have adapted to use the introduced apple (*Malus domesticus*) as a host during the last 150 years (Bush 1969; Feder et al 1988; Linn et al 2004; Meyers et al 2020). It remains an empirical challenge to resolve to which extent morphological adaptations are parallel and predictable under parallel selection regimes (Bolnick et al 2018). Few studies have explicitly compared parallelism in multiple traits in such scenarios, but a recent study of Bahamas mosquitofish (*Gambusia hubbsi*) suggests that only a few traits are highly predictable across parallel predation environments (Langerhans 2018).

Parallelism in response to co-existence with congeners provide a further empirical challenge, as evolutionary divergence may be affected by the local presence or absence of the other incipient species (Amarasekare 2003; Calabrese and Pfennig 2020). Reducing gene flow to populations adapted to other niches is crucial for ecological specialization to result in diversification, as ecological divergence does not necessarily lead to speciation (Taylor et al 2006; Bolnick 2011; Lackey and Boughman 2017). Divergence and specialization may be reversed in the absence of reproductive isolation (Taylor et al 2006; Lackey and Boughman 2017). Hence, if reproductive isolation between the incipient species is incomplete, secondary contact may break up important ecological adaptations. This could be particularly evident when different populations of the diverging species are inhabiting discrete niches, like in the case of insect host races. Close congeners inhabiting different niches may be poorly adapted to the alternative environment (Rundle and Nosil 2005; Nilsson et al 2017; Cronemberger et al 2020; Martin et al 2020), which may lead to reinforcement (Servedio and Noor 2003) and character displacement with stronger differences in traits involved in mate choice (Hinojosa et al 2020). While parallel divergence is expected for ecologically selected traits (Fig. 1a), either parallel- or nonparallel character displacement (Fig. 1b,c) or reduced divergence due to introgression (Fig. 1d) could be expected for traits that are not strongly coupled with host plant use.

**Figure 1.**
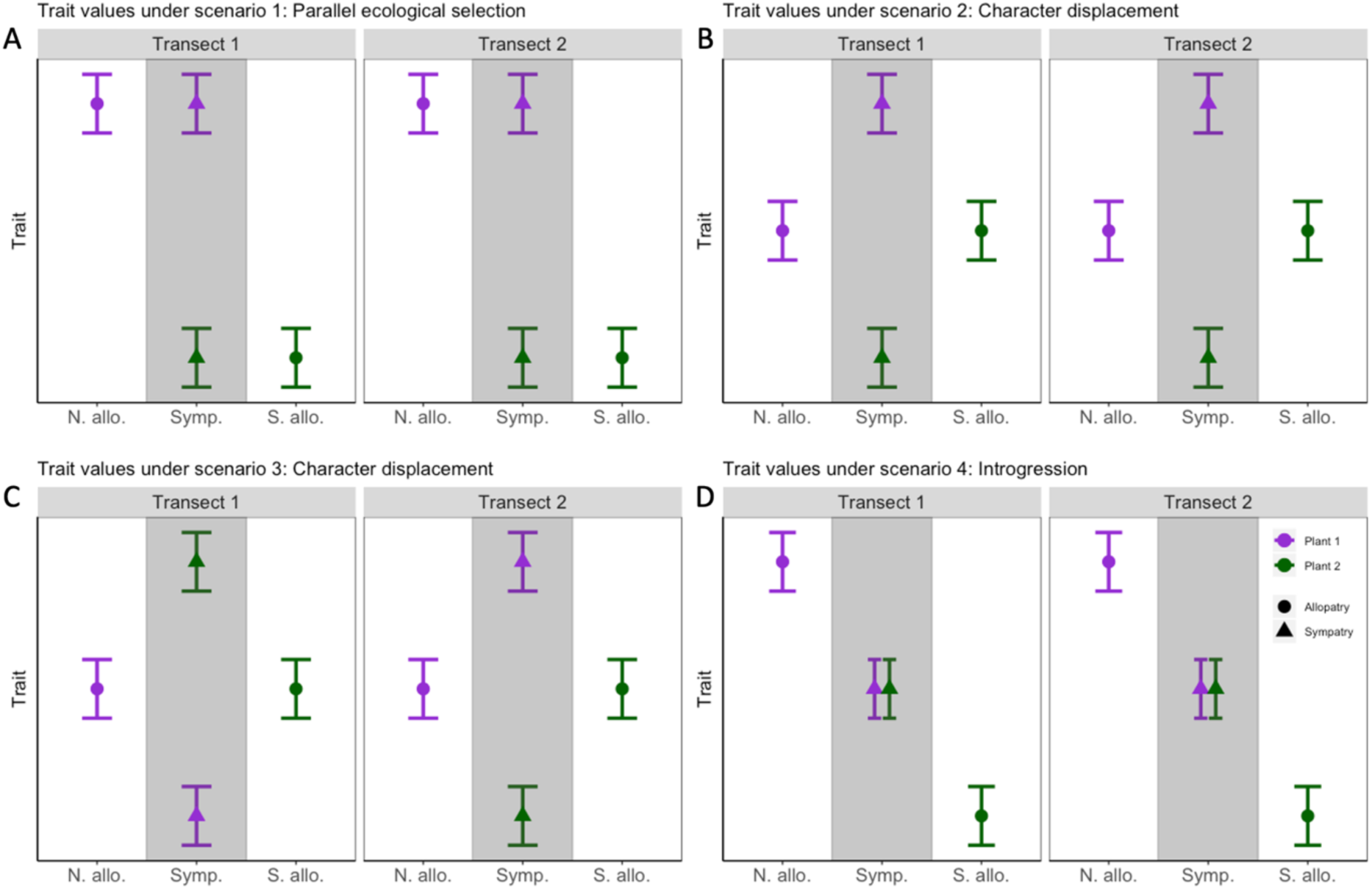
Predictions for morphological divergence under different scenarios. Four possible scenarios for population differentiation depending on the ecological and/or sexual selection pressures at play. Bars represent mean values of morphological traits, with one population being represented by each bar. While the white areas of the graph depict allopatric populations, the grey area includes co-existing populations. ‘N. allo.’ stands for northern allopatry, ‘Symp.’ stands for sympatry and ‘S. allo.’ stands for southern allopatry. Two hypothetical transects are depicted side by side. **A:** Scenario 1, parallel ecological selection between host races. This scenario would be expected for ecologically important traits where natural selection imposed by the environment provided by the host differ. **B**: Scenario 2, character displacement in a shared direction across transects. Should a sexual character be reinforced by maladaptive hybridization consistently across transects, this scenario where traits involved in mate recognition diverge further in sympatry due to character displacement would be expected. **C**: Scenario 3, character displacement in different directions across transects. This would be expected in a sexual character which is reinforced by maladaptive hybridization, similarly to Scenario 2, however in this scenario the direction of adaptation is arbitrary, which would be expected under a mutation order regime. **D:** Scenario 4, hybridization in sympatric areas is causing introgression resulting in reduced divergence in these areas.

To understand whether selection to reduce maladaptive mating in the presence of conspecifics results in parallel morphological divergence to the same extent as ecological adaptations to a novel niche, it is necessary to compare patterns of divergence in allopatry and sympatry across geographic regions. Such studies would test the hypothesis that morphological responses to sympatry can be more arbitrary than adaptations of ecological traits, which are expected to diverge in parallel across populations of similar niches (Fig. 1). In particular, studies asking how co-existence with congeners affects trait distributions, e.g. through parallel- or non-parallel character displacement (Fig. 1), may provide insight into the nature of the selection pressures exerted by co-existence of close congeners. Here, we address how morphological traits evolve and diversify in allopatric and sympatric areas using two host races of the fly *Tephritis conura*. This species has recently undergone a host shift resulting in host races specializing on utilizing different *Cirsium* thistle species as larval food plants (Diegisser et al 2006a; Diegisser et al 2006b; Diegisser et al 2007; Diegisser et al 2008).

Macroevolutionary analyses reveal that host plant driven speciation, as in the case of *T. conura*, is one of the main factors explaining the tremendous diversity of phytophagous insects as many speciation events can be attributed to host plant shifts (Berlocher and Feder 2002; Dres and Mallet 2002; Nylin et al 2014). Moreover, diversification rates are elevated in herbivorous insects compared to their non-herbivorous relatives (Mitter et al 1988; Farrell 1998; Wiens et al 2015). For an insect specialist like *T. conura*, the host plant provides a discrete environment imposing a multidimensional selection pressure. Hence, insect host plant adaptations are predicted to affect multiple traits involved in e.g. phenological matching, female host preference and larval performance (Matsubayashi et al 2010). The putatively strong selection pressures involved in host plant adaptation provide a strong prediction of parallelism in host-use related traits of phytophagous insects at early stages of the speciation process (Nosil et al 2002; Meyers et al 2020). However, the prediction is less straightforward for responses to sympatry, because the traits involved in putative reinforcement are not necessarily associated with host use.

*Tephritis conura* provides an excellent system for studying parallelism in divergence in response to ecological adaptation and co-existence with congeners, as it enables comparisons of trait variation across replicated sympatric and allopatric settings. The two recently established host races inhabit two geographically isolated sympatric zones with adjacent allopatric populations, one East and one West of the Baltic Sea (Fig. 2a). *Tephritis conura* females of the two host races are ovipositing into buds of either *C. heterophyllum* (the ancestral host; Fig. 2b) or *C. oleraceum* (the novel host; Fig. 2c), and larvae show host specific performance and survival (Diegisser et al 2008). Hence, the system enables investigating response to sympatry across two geographically separated contact zones. Specifically, we test the hypotheses (i) that there are consistent morphological responses to a host plant shift across the Eastern and Western areas and (ii) that responses to sympatry are parallel in the two contact zones. Given the consistent genetic differentiation between host races both in allopatry and sympatry (Diegisser et al 2006a; Ortega et al *in prep*) we predict the host races to be morphologically divergent, and this divergence to be consistent across geographical settings. As larval survival is strongly reduced in the alternate host (Diegisser et al 2008), we predict hybridization to be maladaptive, potentially resulting in reinforcement and character displacement between the host specialists in sympatry.

**Figure 2:**
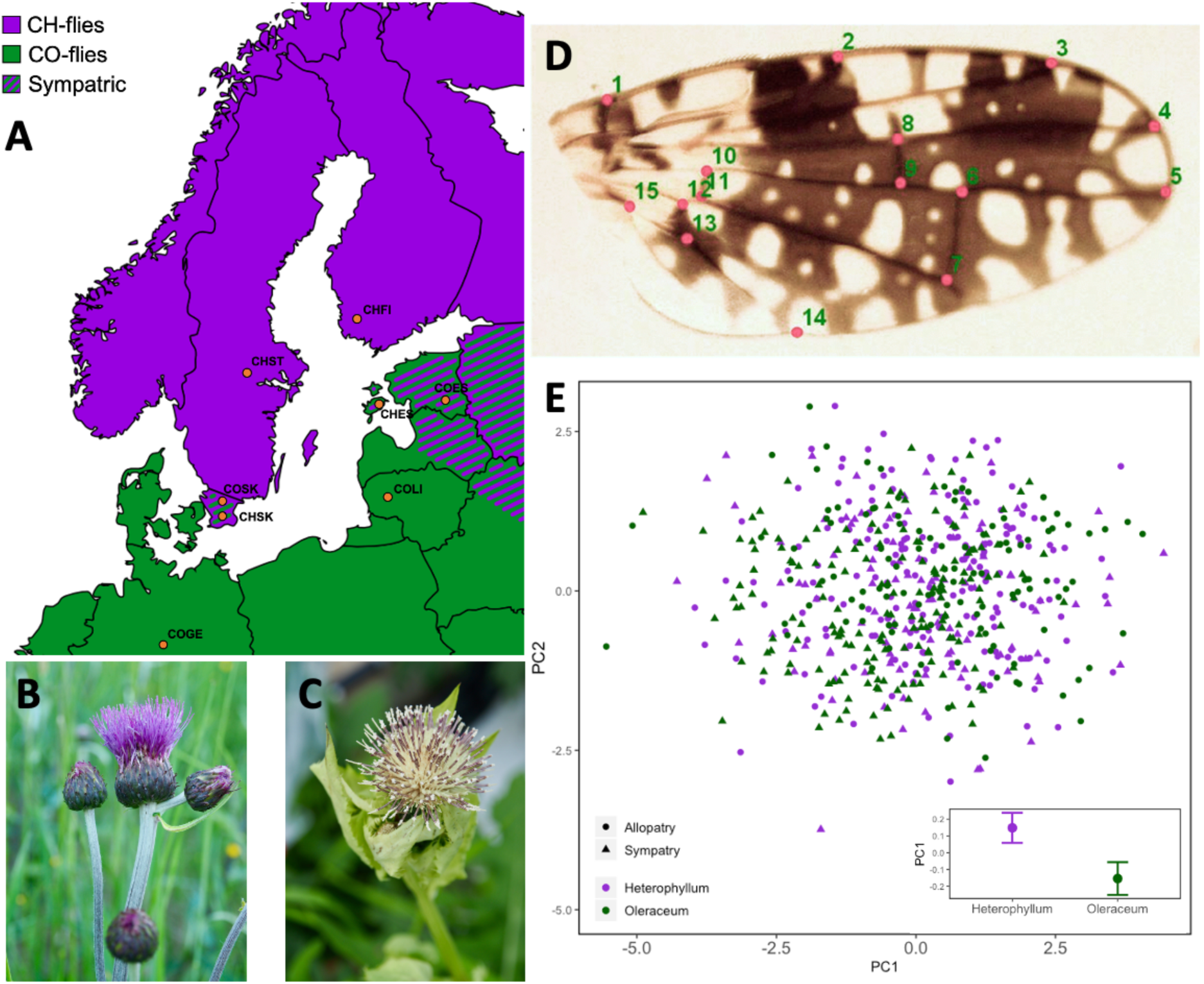
Methods. **A**: Map showing distribution of the *T. conura* host races, with the distribution of CH-flies infesting *Cirsium heterophyllum* depicted in purple and that of CO-flies infesting *Cirsium oleraceum* depicted in green. Sampling locations are indicated by red circles. **B**: *Cirsium heterophyllum*. **C**: *Cirsium oleraceum*. **D**: Dorsal photograph of a *T. conura* wing annotated with 15 landmarks used to analyze wing shape. **E**: Principal component analysis of *T. conura* morphology based on females. The first two principal components explain 37.84% of the variation. Insert shows the mean and standard error of the mean of PC1 in the two host races.

## Methods

### Study species and sampling

The dipteran *T. conura* infest several species of the thistle genus *Cirsium* (Asteraceae) (Romstock-Volkl 1997). Adult *T. conura* oviposit into thistle buds during early summer, wherein the adolescent flies remain during larval development and pupation. *Tephritis conura* infesting *Cirsium heterophyllum*, the melancholy thistle, have recently colonized and adapted to *Cirsium oleraceum*, the cabbage thistle. Haplotype analyses suggest that a peripatric host shift took place during the last ice age in the Alps (Diegisser et al 2006b). The two host races are largely reproductively isolated, but there is evidence of some small amounts of gene flow (Diegisser et al 2006b). The flies infesting *C. oleraceum* have adapted ecologically to the smaller bud sizes (Diegisser et al 2007), and have a significantly shorter ovipositor to body size ratio. Another variable character is wing pigmentation. Wing patterns in Tephritid flies have been suggested to be under sexual selection, as males perform dances with their wings to attract females (Sivinski and Pereira 2005). Tephritid males attempting to initiate copulation situate themselves in front of female flies and posture with their wings (Sivinski 2000; but see Briceno and Eberhard 2017). We therefore have reason to assume that ecological selection pressures are more important for size traits in general, and for ovipositor length in particular, while wing traits are more likely shaped by sexual selection. For simplicity, we refer to flies infecting *C. heterophyllum* as CH-flies, while the flies infecting *C. oleraceum* will be denoted CO-flies.

We used a parallel sampling design to examine phenotypic adaptation to a novel host plant and effects of co-existence between host races, by sampling each host race both in sympatry and allopatry on each side of the Baltic (Fig. 2a). We collected thistle buds infested by *T. conura* larvae/pupae and allowed the adults to eclose in a common environment. CO-fly larvae were sampled in Germany and Lithuania (allopatric areas), and both host races were collected in sympatric areas in southern Sweden and Estonia. Allopatric CH-fly larvae were sampled in central-Sweden and Finland (Fig. 2a, Table S1). All sampling took place during June and July 2018. The sampling scheme enables examining to what extent patterns of phenotypic divergence are explained by host plant adaptations, by co-existence with the other host race, and if these patterns differ between the two transects. Typically, *C. heterophyllum* and *C. oleraceum* do not grow in the same microhabitat. Thus, the sympatric and allopatric definitions here refer to the presence of one or both thistle species in a region (Fig. 2a).

### Morphological measurements

*Tephritis conura* adults eclosed from field-collected thistle buds in a common lab environment, (see Supplementary material S1). One male and one female per bud were euthanized by freezing a few days after emergence and subsequently included in the morphological analysis. For each individual, we took magnified photographs using a Celestron 44308 USB microscope. We photographed a lateral image of the fly body after removal of the wings and a dorsal image of the right wing on a transparent background to allow better visibility of the wing veins. Body length and ovipositor length were measured digitally from lateral photographs (Fig. S1). We placed 14 landmarks, adapted from Pieterse et al. (2017), digitally on the dorsal wings (Fig. 2d,S2) for geometric morphometrics (Zelditch 2004). We added a landmark; number 15, to reflect the high variance in the proximal area on the wing. Digitization was performed in TPSDig2 v2.31(Rohlf 2017) and we used TPSUtil v1.76 (Rohlf 2018) for file handling. Wing melanisation was measured with an automated script without any user queries developed in MATLAB (Matlab 2017). As the wings do not display any chromaticity, analysis is based on the red spectral band only. The script extracted the intensity of the red spectral band for each pixel, and performed white calibration by division with the statistical mode corresponding to the white background of the images. Subsequently, images were inverted to represent absorbance rather than reflectance, so the melanised wing would extend from the white background (set to zero). The script identified the wing and separated melanised areas from non-melanised areas, and the size of these areas were divided to estimate the fraction of the wing that was melanised.

### Statistical analysis

We used PAST3 v3.20 (Hammer et al 2001) to apply a Procrustes fit to the landmark data to align and scale the wings (Fig. 2d). To produce relative warps (i.e. principal components of shape) to compare shape between groups, a wing-shape principal component analysis (PCA) was performed with the Procrustes fitted data using PAST3 v3.20 (Hammer et al 2001) (Fig. S3). Based on the variance explained by the eigenvalues (Fig. S4) and the broken stick criterion (Jackson 1993), six relative warps, principal components of shape, which jointly explain 68% of the wing shape variance were identified (Fig. S5). These relative warps were included in subsequent analyses of phenotypic divergence to represent wing shapes. All subsequent statistical analysis were performed in the statistical software R (R Core Team 2019).

We quantified five morphological traits (body length, ovipositor length, wing length, wing width and melanisation ratio) in addition to wing shape (represented by relative warps produced from landmark analysis) for 583 flies. As an exploratory analysis to investigate whether host race was the major factor explaining variation in fly morphology, we performed a full PCA on the variables fly body length (mm), wing length (mm), wing width (mm), wing melanisation ratio (%) and relative warps 1-6 reflecting wing shape, excluding ovipositor length to be able to include both sexes. We identified four significant dimensions of variation from the full PCA analysis using the broken stick criterion (Jackson 1993) (Fig. S6). Collectively, these full PCA-axes explained 78% of the morphological variance in the dataset.

To formally test if the two host races were significantly differentiated, we applied a multivariate analysis of variance (MANOVA), with all variables measured included as response variables (body length, ovipositor length, wing width, wind length, wing melanisation and PC1-6 of wing shapes). To further test if the patterns of morphological adaptation were parallel in the Eastern and the Western transects, and explicitly address if co-existence affected host plant races in the same way in these replicates, we performed a full MANOVA with host race, co-existence and geographical setting and their 3-way and 2-way interactions as factors. The MANOVAs were performed separately on females and males as this enabled including the biologically important trait ovipositor length (Diegisser et al 2007) in analyses of females. This division of sexes reduced the multicollinearity of explanatory factors to below recommended values (Hair, 2010).

To further investigate parallelism in differentiation of the host race pairs, we applied separate linear discriminant function analyses (LDAs) on the data from the Eastern and Western transects. We used host races and co-existence with the other host race as factors in the models. This analysis was performed on males and females separately. To test if the patterns of divergence differed significantly between transects, we performed each LDA 10,000 times using the bootstrap R package ‘boot’ (Canty and Ripley 2020) and used the confidence intervals to assess if the loadings differed between analyses. In addition, we assessed the proportions of divergence that is shared among host races and unique among the populations, respectively, in a nested MANOVA using size and shape variables against host race and transect following Langerhans & DeWitt (2004).

## Results

We found no strong evidence for separation between host races along the two major axes of divergence when including both sexes (Fig. 2e). CO-flies had, on average, lower values of PC1 compared to CH-flies, reflecting smaller body and wing size, as predominantly size variables load positively on PC1 (GLM: F_1, 581_ = 5.189, p = 0.023; Fig. 2e; Table S2). The significant host race differences also hold true in a more complex model, including co-existence and geographic origin (see below; Table 1). However, the difference between host races was small compared with the variation among populations within host races. There was no separation between host races along the second PC axis, with higher values representing higher wing melanisation and lower representation of wings that are larger than average in the distal and proximal areas of the wing, while being smaller in the posterior area, see relative warp 5 in Fig. S5.

**Table 1:** MANOVA. Addressing the effect of fly host race, co-existence, transect and the interaction of these factors on multivariate morphological divergence in five morphological and six wing shape traits for females and four morphological traits and six wing shapes for males.

| MANOVA | df | Pillai | Approx. F | num df | den df | P |  |
| --- | --- | --- | --- | --- | --- | --- | --- |
| Females |  |  |  |  |  |  |  |
| Host plant | 1 | 0.258 | 8.42 | 11 | 267 | < 0.001 | *** |
| Co-existence | 1 | 0.295 | 10.15 | 11 | 267 | < 0.001 | *** |
| Baltic Sea | 1 | 0.374 | 14.52 | 11 | 267 | < 0.001 | *** |
| Host plant x Co-existence | 1 | 0.218 | 6.79 | 11 | 267 | < 0.001 | *** |
| Host plant x Baltic Sea | 1 | 0.18 | 5.31 | 11 | 267 | < 0.001 | *** |
| Co-existence x Baltic Sea | 1 | 0.163 | 4.74 | 11 | 267 | < 0.001 | *** |
| Host plant x Co-existence x Baltic Sea | 1 | 0.117 | 3.22 | 11 | 267 | < 0.001 | *** |
| Residuals | 277 |  |  |  |  |  |  |
| Males |  |  |  |  |  |  |  |
| Host plant | 1 | 0.078 | 2.29 | 10 | 271 | 0.014 | * |
| Co-existence | 1 | 0.194 | 6.54 | 10 | 271 | < 0.001 | *** |
| Baltic Sea | 1 | 0.393 | 17.51 | 10 | 271 | < 0.001 | *** |
| Host plant x Co-existence | 1 | 0.179 | 5.92 | 10 | 271 | < 0.001 | *** |
| Host plant x Baltic Sea | 1 | 0.16 | 5.12 | 10 | 271 | < 0.001 | *** |
| Co-existence x Baltic Sea | 1 | 0.131 | 4.1 | 10 | 271 | < 0.001 | *** |
| Host plant x Co-existence x Baltic Sea | 1 | 0.092 | 2.74 | 10 | 271 | 0.003 | ** |
| Residuals | 280 |  |  |  |  |  |  |

Intriguingly, host plant and co-existence affected female morphology differently East and West of the Baltic Sea, as illustrated by a significant 3-way interaction (Pillai’s trace = 0.117, *F*_11, 277_ = 3.22, p <0.001; Table 1). All main effects and 2-way interactions were also significant in the full model (Table 1). These patterns hold true also for males (Pillai’s trace = 0.092, *F*_10, 271_ = 2.74, p = 0.003; Table 1). Hence, depending on geography, host race and co-existence affected fly morphology differently. Co-existence with the other host race decreased morphological divergence in both females and males (Pillai’s trace = 0.295, *F*_11, 277_ = 10.15, p <0.001 and Pillai’s trace = 0.194, *F*_10, 271_ = 6.54, p <0.001 respectively), and flies from the two transects differed significantly in morphology for both sexes (Pillai’s trace = 0.374, *F*_10, 271_ = 14.52, p <0.001 for females and Pillai’s trace = 0.393, *F*_10, 271_ = 17.51, p <0.001 for males).

Interestingly, host race differences in females depend on co-existence, with CO-fly females becoming more similar in sympatry compared to CH-fly females (Pillai’s trace = 0.218, *F*_11, 277_ = 6.79, p <0.001; Tables 1 and S3; Fig. 3 and S7). In contrast to females, co-existence affected CO-fly males more strongly, with CO-fly males becoming more similar to CH-fly males in sympatry whereas CH-fly males were more similar across all populations (Pillai’s trace = 0.179, *F*_10, 271_ = 5.92, p <0.001; Tables 1 and S3; Fig. S8-S11). The differences between host races varied between transects in females, with a stronger host race divergence in the Western than the Eastern transect (Pillai’s trace = 0.18, *F*_11, 277_ = 6.79, p <0.001; Tables 1 and S3; Fig. 3 and S7; Tables 1 and S3; Fig. 3 and S7). This pattern holds true also for males (Pillai’s trace = 0.16, *F*_10, 271_ = 5.12, p <0.001; Tables 1 and S3; Fig. S8-S11). Finally, the differences between allopatric and sympatric populations are stronger in Western flies compared to Eastern both in females (Pillai’s trace = 0.163, *F*_11, 277_ = 4.74, p <0.001; Tables 1 and S3; Fig. 3 and S7) and males (Pillai’s trace = 0.131, *F*_10, 271_ = 4.1, p <0.001; Tables 1 and S3; Fig. S8-S11).

**Figure 3:**
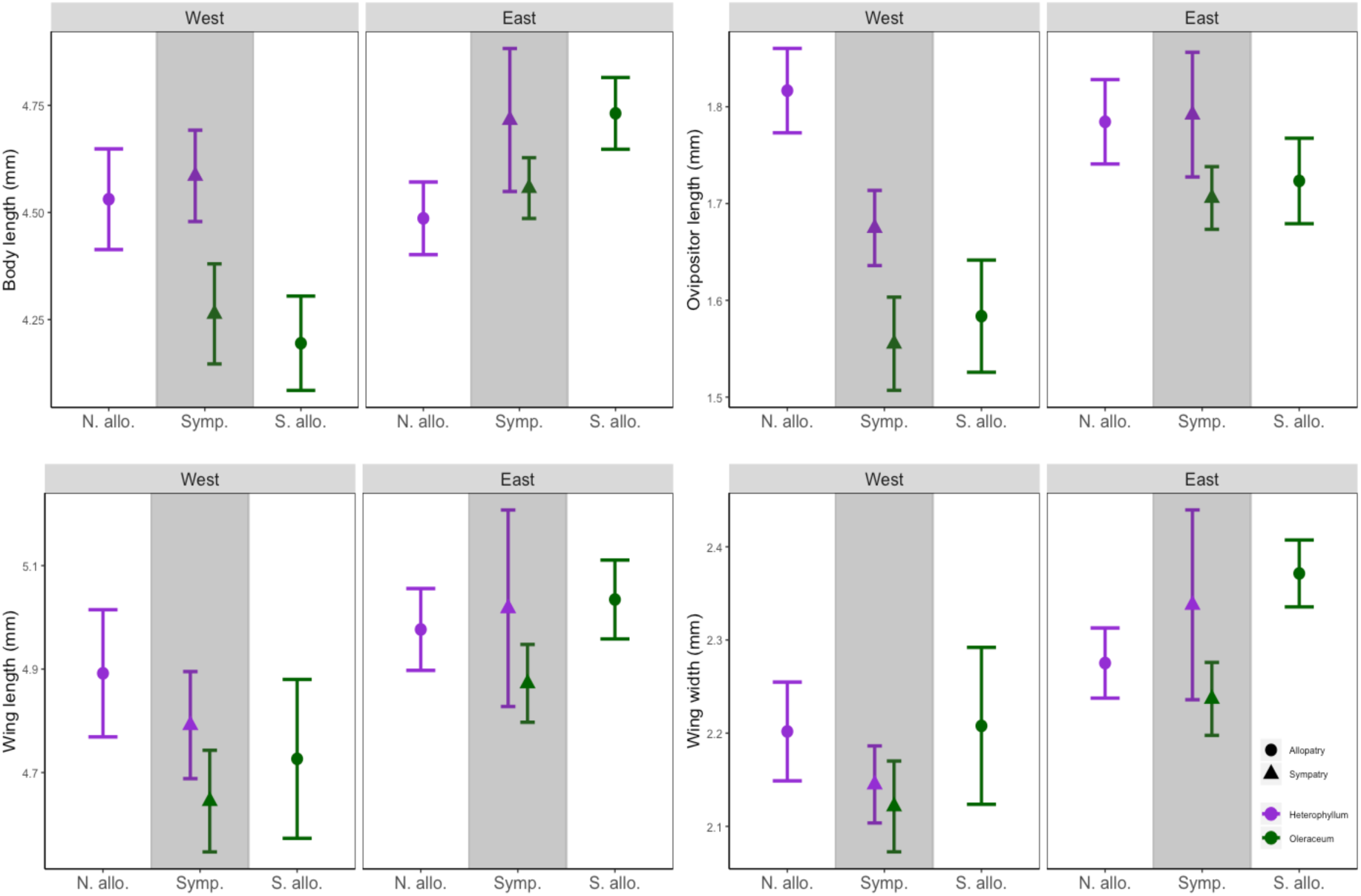
Trait measurements of female *T. conura* divided by host race, co-existence and geographic setting. Host race is illustrated by color, with purple corresponding to CH-flies infesting purple melancholy thistle and green to CO-flies infesting white cabbage thistle. The grey shaded area in the center contains the sympatric populations (denoted by triangles). X axis labels represent which state of co-existence the fly population is in. ‘N.allo.’ stands for Northern allopatric, ‘Symp.’ stands for sympatric and ‘S. allo.’ stands for Southern allopatric. West and East headers represent from which side of the Baltic Sea the populations are sampled. **A**: Mean values of female *T. conura* body length per population. **B:** Mean values of female *T. conura* ovipositor length per population. **C:** Mean values of female *T. conura* wing length per population. **D:** Mean values of female *T. conura* wing width per population. All plots portray mean trait values with 95% confidence interval bars of the mean.

To assess how much of the female divergence was unique and shared between host races we also estimated Wilk’s partial η^2^. While a high share of the partial variance was explained by shared divergence between host races (*F*_11.273_ = 4.96, 30.6%) and transect specific patterns of host race divergence (*F*_11.273_ = 3.81, 25.3%), divergence between transects explained the highest percentage of partial variance (*F*_11.273_ = 6.85, 37.8% (Table S4).

As host race affected morphology differently depending on geography, we further tested if the major axis of divergence separated host races in both transects, and if the same traits separated groups using a Linear Discriminant Analysis (LDA). This analytical approach revealed that the importance of host race for population separation differed between transects. In the Western LDA, host races separated along the first discriminant axis whereas the sympatric and allopatric populations separated along the second discriminant axis (Fig. 4). In the Eastern LDA, the first discriminant instead divided the two CO-fly populations and the second discriminant axis divided the two CH-fly host race populations (Fig. 4). These analyses also revealed that the host races from transects East and West of the Baltic had diverged in different sets of characters as bootstrap loadings from the two LDAs show that different characters loaded on the first discriminant axes (Table S5). For the Eastern transect, body length and wing warp 6 had positive loadings, and wing size, ovipositor length and wing warp 1 had negative loadings on LD1, whereas for the Western transect, LD1 had positive loadings for wing width and shape. In the Eastern transect, wing shape, body length and wing width had negative loadings, but wing length positive loadings on LD2, while for the Western transect body length, ovipositor length and wing lengths loaded positively on LD2. Wing warp 5 loaded positively on LD1 for both transects, but the loading was substantially stronger for Western populations.

**Figure 4:**
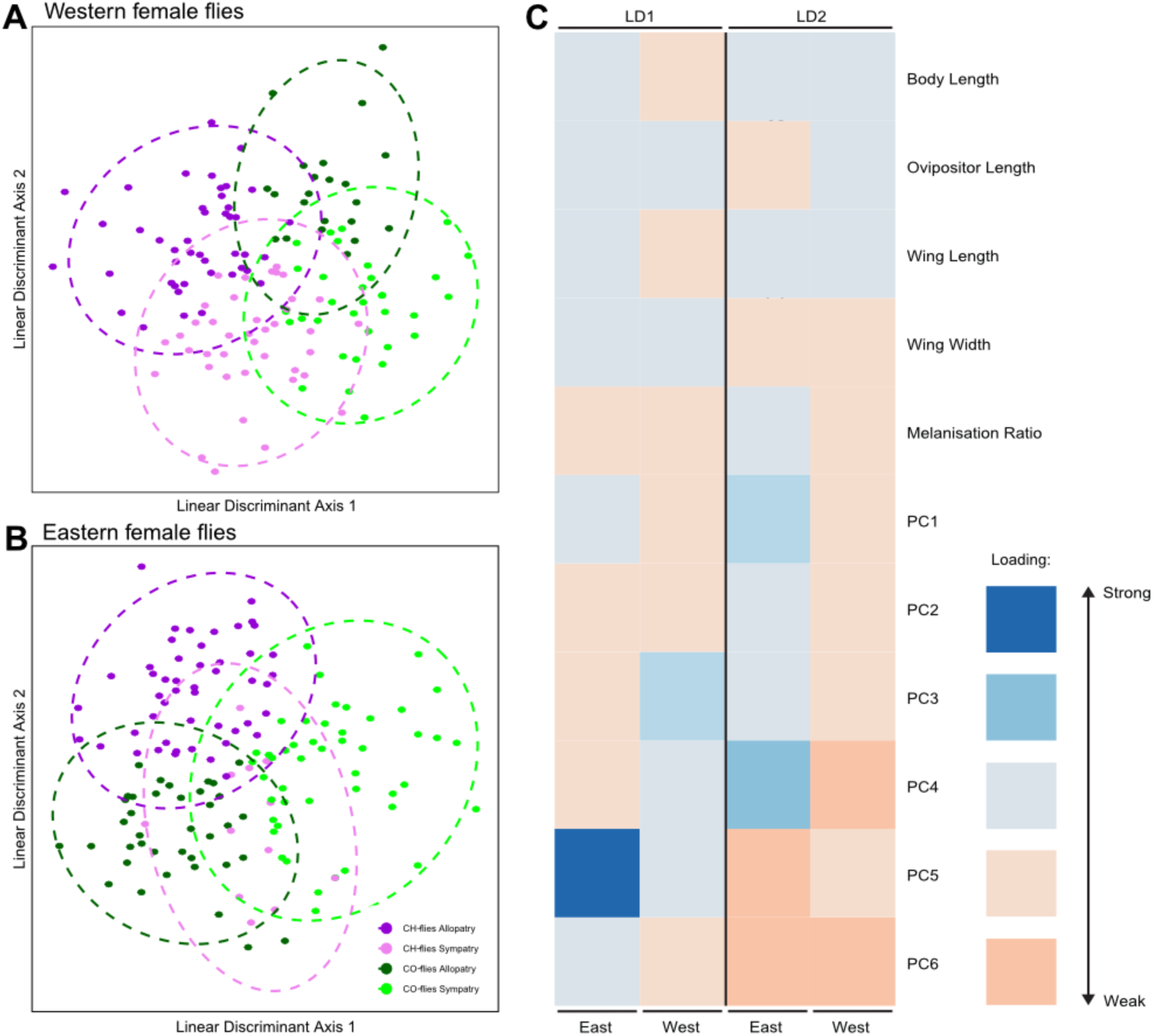
Linear discriminant function analyses and bootstrapped loadings. **A and B**: LDAs illustrating differences in how the host races group along the first two linear discriminant axes in Western (A) and Eastern (B) flies. **C:** The morphological traits loading on the two first discriminant axes for Eastern and Western flies. Colors illustrate how much the standard error diverges from zero based on 100 000 bootstrap replications. Loading that surpass zero are depicted in red colors whereas loadings significantly lower than zero are colored in blue. Plots and analyses are based on female fly morphology.

## Discussion

Contrary to previous studies that found clear differences between the *T. conura* host races (Diegisser et al 2007) we found low and variable morphological differentiation. The multivariate differentiation between host races was driven by subtle differences across many traits, and the traits that were divergent between host races differed in the two transects. Host races separated slightly but significantly along the main axis of variation, reflecting mainly ovipositor length differences, with CH-fly ovipositors being slightly longer than CO-fly ovipositors, but with unexpected substantial variation among populations. Previous work on the *T. conura* host races has reported consistent divergence for loci inferred to be involved in host plant adaptation (*Peptidase D* and *Hexokinase*; see (Diegisser et al 2006b), and recently obtained whole genome data (Ortega et al *in prep*) support the presence of discrete genetic clusters for each host race. Moreover, the flies also have poor performance on the alternate host plant (Diegisser et al 2008). In light of this, the utterly moderate and variable host race divergence in morphology was unexpected. However, our findings are in line with the observation that parallelism in fitness is typically higher than parallelism in both phenotypic divergence and genetic divergence (Bolnick et al 2018). Patterns of parallel divergence only in traits under very strong ecological selection is consistent with the findings of parallelism in response to different predation regimes in Bahamas Mosquitofish (Langerhans 2018), where only a few traits show highly predictable patterns of diversification. Hence, at early stages of diversification driven by ecological adaptation, parallelism may be high only for traits that are strongly coupled to the ecological factor that adaptively diversify the incipient species.

We find that two out of five female traits, ovipositor length and body length, match a parallel divergence scenario, and body length only in the Western transect (Fig. 1a). Hence, the degree of parallelism among host races in morphological divergence was lower than expected. Ovipositor length was significantly diverged between host races, but the strength of divergence differed between sympatric and allopatric populations as well as between the Eastern and Western populations. In Western populations, the sympatric CH-fly population has intermediate ovipositor lengths, whereas CO-flies and Eastern CH-flies show parallel ecological divergence between sympatry and allopatry, corresponding to the expectations under parallel ecological selection (Fig. 1a). The intermediate measurements for traits found in the Western CH-flies could suggest selection for shorter ovipositors in sympatry, or reflect introgression. We find host race divergence in body size, with clear host race differences in body length in the Western transect (corresponding to Fig. 1a), while Eastern flies have a shuffled distribution of body sizes. Size typically varies with temperature in insects (Atkinson 1994; but see Shelomi 2012). While the parallel sampling design would partly correct for temperature effects, temperature could still differ between the Eastern and Western transects.

When we jointly investigated effects of co-existence, geographic origin and their interactions on host race divergence, we found that both co-existence with the other host race and whether the population originate from East or West of the Baltic Sea affected morphology (Fig. 3). Hence, the low host race divergence may, to a high extent, be explained by interacting effects of co-existence with the other host race, with non-parallel patterns of divergence in the two transects, which could indicate a mutation order scenario (Fig. 1c) (Mendelson et al 2014). If reinforcement due to maladaptive hybridization would have been a prominent force acting on the *T. conura* host races, we would expect traits to be more divergent between host races in sympatry compared to allopatry, i.e. for character displacement to arise (Comeault et al 2016; Calabrese and Pfennig 2020; Kyogoku and Wheatcroft 2020; Fig. 1b,c). This was generally not the case. Instead, the most common pattern was that traits in sympatric populations are more similar between host races than traits in allopatry. Contrary to our expectations, based on the fact that *Tephritid* flies perform a mating ritual that includes elaborate wing movements from both sexes during courtship (Sivinski 2000; but see Briceno and Eberhard 2017), this pattern was strongest for wing morphology traits. This is the opposite of what would be predicted under a reinforcement scenario but consistent with introgression (Fig. 1d), as found by Ortega et al. [*in prep*.], or effects of a shared environment.

Wing size was most strongly affected by co-existence. Interestingly, wing length was overall significantly shorter in sympatry than in allopatry, consistent on both sides of the Baltic Sea. Disproportionally shorter wing length in relation to body size could be consistent with selection for shorter dispersal distances (Claramunt et al 2012), potentially to avoid dispersal to the alternate host plant which could lead to maladaptive introgression. We also found patterns consistent with character displacement, as sympatric flies have similarly narrower wings than their respective allopatric counterparts in the Western transect. In contrast, Eastern sympatric CH-flies have broader and sympatric CO-flies narrower wings. These observations are consistent with a mutation order scenario, where the difference may be important but the specific trait or direction of the difference is arbitrary (Mendelson et al 2014). Some degree of sexual selection may further differentiate populations under a mutation order scenario (Rundle and Rowe 2018), underlining the need to investigate if there is sexual selection at work in this system and disentangling how it would operate.

Contrary to studies of parallelism in sexually selected traits in ecotypes in e.g. sticklebacks (Boughman et al 2005), we find mixed evidence for parallelism. While shared host race divergence constituted the main share of variation, and host races are separated by the main discriminant axis in the Western transect, host race was a less important predictor of divergence in the Eastern transect where the two CO-fly populations instead separate along the first discriminant axis. The traits that differ between host races also differs between our two geographic replicates. The differences in patterns between the transects East and West of the Baltic Sea could have several additional or alternative explanations related to demographic history, population size and the extent to which the host plant races co-exist locally. Possibly, one contact zone may be older than the other and populations in older sympatry would have had more time for character displacement to develop. Alternatively, if the Eastern transect has a higher proportion suitable thistle habitat, this could have increased both within- and between host race connectivity and potentially gene flow (Servedio and Noor 2003). Genetic data and detailed analyses of introgression should be used to resolve whether selection against hybridization could be expected.

Morphological differentiation does not always strongly reflect even crucial ecological adaptations. For instance, cultural evolution contributes to reproductive isolation in Cassia crossbills, *Loxia* sinesciuris (Porter and Benkman 2019) and *Rhagoletis pomonella* have adapted their phenologies to host fruits ripening at different times of the year (Filchak et al 2000). Another potential explanation to the low consistency of host race divergence may be the traits included in this analysis. Females were more diverged than males likely as ovipositor length was differentiated, consistent with previous findings (Diegisser et al 2007). These findings are similar to those of Jourdan et al. (2016) where divergence in female mosquitofishes (*Gambusia*) divergence was more parallel than male divergence. Potentially, the other traits measured are not important enough for host plant adaptation to result in strongly parallel divergence.

Traits that have been shown to differ strongly between host races include female ovipositor length (Diegisser et al 2007) and the larval ability to survive on the different host plant species (Diegisser et al 2008). A performance experiment showed low viability of *T. conura* larvae adapted to *C. heterophyllum* when reared on *C. oleraceum* (Diegisser et al 2008). Thus, the larval ability to process plant tissue is likely under strong selection, and expected to show similar patterns across populations. Furthermore, host plant preference may have a potential to act as a magic trait (Gavrilets 2004; Thibert-Plante and Gavrilets 2013) in *T. conura*, separating the habitats of the populations and simultaneously providing reproductive isolation as these flies mate on their host plant (Diegisser et al 2007). Finally, we cannot rule out that other, unmeasured traits are important for sexual selection, as pheromones have been suggested to play a role in tephritid mate choice (Roriz et al 2019), as well as overexpression of antioxidants, which have been shown to increase male performance under certain conditions (Teets et al 2019). Regardless of the exact selection pressures acting on our set of study traits, our findings are important, because they show a context dependence for host race adaptation. Parallelism was found only in ecologically strongly selected traits, whereas we found non-parallel changes, some consistent with character displacement for other traits. These insights should guide the design and interpretation when studying ecologically driven divergence.

In conclusion, our work suggests that morphological responses to niche shifts can be highly context dependent. Co-existence with closely related congeners and demographic origin may affect easily measured morphological characters, potentially masking underlying parallelism in traits important for adaptation to specific niches. Moreover, we find intriguing patterns of non-parallel divergence to co-existence in the two geographic replicas, suggesting that mutation order dependent divergence potentially could lead to different solutions to avoid introgression in independent contact zones.

## Supporting information

All supplementary material

## Acknowledgements

We thank Mikkel Brydegaard for help with automatizing wing color analysis, Jes Johannesen for helpful advice on *Tephritis conura* ecology, Øystein Opedal for helpful statistical discussions, Jodie Lilley, Emma Kärrnäs and Mathilde Schnuriger for help during field- and lab work, and the members of the Lund University research group on the Evolutionary Ecology of Plant-Insect Interactions for valuable discussions on earlier drafts of this manuscript. This study was financed by a Wenner-Gren Fellowship, a Crafoord grant and a Swedish Research Council grant to AR.

## Declarations

### Funding

This study was financed by a Wenner-Gren Fellowship, a Crafoord grant and a Swedish Research Council grant to AR.

### Conflicts of interest

The authors declare no conflicts of interest.

### Ethics approval

Not applicable

### Consent to participate

All co-authors consent to participation.

### Consent for publication

All co-authors consent to publication.

### Availability if data and material

All data will be deposited in dryad (upon acceptance of the MS).

### Code availability

The code used to analyse data is available at github.com/kalnil/tephritis.

### Author contribution

A.R conceived and designed the study. KJ.N., J.O. and A.R. performed the field work. KJ.N. reared the flies, quantified the morphological traits and performed the statistical analyses, with advice from A.R. and M.F. KJ.N. wrote a draft of the manuscript. A.R. and M.F. helped writing the manuscript and all co-authors commented on and approved the final version of the manuscript.

## Notes

### Competing Interest Statement

The authors have declared no competing interest.

## References

Amarasekare P (2003) Competitive coexistence in spatially structured environments: A synthesis. Ecology Letters 6:1109–1122

Atkinson D (1994) Temperature and organism size - A biological law for ectotherms. In: Begon M & Fitter AH (eds) Advances in Ecological Research, vol 25. Academic Press Ltd-Elsevier Science Ltd, London, pp1–58

Baldwin BG, Carlquist S & Carr GD 2003 Tarweeds & silverswords: evolution of the Madiinae (Asteraceae). Missouri Botanical Garden Press, St. Louis, Mo.

Bay RA, Arnegard ME, Conte GL, Best J, Bedford NL, McCann SR, Dubin ME, Chan YF, Jones FC, Kingsley DM, Schluter D & Peichel CL (2017) Genetic coupling of female mate choice with polygenic ecological divergence facilitates stickleback speciation. Current Biology 27:3344–3349

Berlocher SH & Feder JL (2002) Sympatric speciation in phytophagous insects: Moving beyond controversy? Annual Review of Entomology 47:773–815

Bolnick DI (2011) Sympatric speciation in threespine stickleback: Why not? International Journal of Ecology 2011:e942847

Bolnick DI, Barrett RDH, Oke KB, Rennison DJ & Stuart YE (2018) (Non)Parallel evolution. Annual Review of Ecology, Evolution, and Systematics 49:303–330

Boughman JW, Rundle HD & Schluter D (2005) Parallel evolution of sexual isolation in sticklebacks. Evolution 59:361–373

Briceno RD & Eberhard WG (2017) Copulatory dialogues between male and female tsetse flies (Diptera: Muscidae: Glossina pallidipes). Journal of Insect Behavior 30:394–408

Bush GL (1969) Sympatric host race formation and speciation in frugivorious flies of genus Rhagoletis (Diptera, Tephritidae). Evolution 23:237–251

Calabrese GM & Pfennig KS (2020) Reinforcement and the proliferation of species. Journal of Heredity 111:138–146

Canty A & Ripley BD (2020) boot: Bootstrap R (S-Plus) functions. R package version 1.3-25.

Claramunt S, Derryberry EP, Remsen JV, Jr. & Brumfield RT (2012) High dispersal ability inhibits speciation in a continental radiation of passerine birds. Proceedings of the Royal Society B-Biological Sciences 279:1567–1574

Comeault AA, Venkat A & Matute DR (2016) Correlated evolution of male and female reproductive traits drive a cascading effect of reinforcement in Drosophila yakuba. Proceedings of the Royal Society B-Biological Sciences 283:20160730

Cronemberger AA, Aleixo A, Mikkelsen EK & Weir JT (2020) Postzygotic isolation drives genomic speciation between highly cryptic Hypocnemis antbirds from Amazonia. Evolution 74:2512–2525

Diegisser T, Johannesen J & Seitz A (2006a) The role of geographic setting on the diversification process among Tephritis conura (Tephritidae) host races. Heredity 96:410–418

Diegisser T, Johannesen J & Seitz A (2008) Performance of host-races of the fruit fly, Tephritis conura on a derived host plant, the cabbage thistle Cirsium oleraceum: Implications for the original host shift. Journal of Insect Science 8:1–6

Diegisser T, Seitz A & Johannesen J (2006b) Phylogeographic patterns of host-race evolution in Tephritis conura (Diptera: Tephritidae). Molecular Ecology 15:681–694

Diegisser T, Seitz A & Johannesen J (2007) Morphological adaptation in host races of Tephritis conura. Entomologia Experimentalis Et Applicata 122:155–164

Dres M & Mallet J (2002) Host races in plant-feeding insects and their importance in sympatric speciation. Philosophical Transactions of the Royal Society B-Biological Sciences 357:471–492

Farrell BD (1998) “Inordinate fondness” explained: Why are there so many beetles? Science 281:555–559

Feder JL, Chilcote CA & Bush GL (1988) Genetic differentiation between sympatric host races of the apple maggot fly Rhagoletis pomonella. Nature 336:61–64

Filchak KE, Roethele JB & Feder JL (2000) Natural selection and sympatric divergence in the apple maggot Rhagoletis pomonella. Nature 407:739–742

Gavrilets S 2004 Fitness landscapes and the origin of species. Princeton University Press, Princeton, N.J.

Hammer Ø, Harper DAT & P. D. R (2001) PAST: Paleontological statistics software package for education and data nalysis. Palaeontologica Electronica 4:9

Hinojosa JC, Koubinova D, Dinca V, Hernandez-Roldan J, Munguira ML, Garcia-Barros E, Vila M, Alvarez N, Mutanen M & Vila R (2020) Rapid colour shift by reproductive character displacement in Cupido butterflies. Molecular Ecology 29:4942–4955

Jackson DA (1993) Stopping rules in principal components-analysis - A comparison of heuristic and statistical approaches. Ecology 74:2204–2214

Jourdan J, Krause ST, Lazar VM, Zimmer C, Sommer-Trembo C, Arias-Rodriguez L, Klaus S, Riesch R & Plath M (2016) Shared and unique patterns of phenotypic diversification along a stream gradient in two congeneric species. Scientific Reports 6:38971

Kyogoku D & Wheatcroft D (2020) Heterospecific mating interactions as an interface between ecology and evolution. Journal of Evolutionary Biology 33:1330–1344

Lackey ACR & Boughman JW (2017) Evolution of reproductive isolation in stickleback fish. Evolution 71:357–372

Langerhans RB (2018) Predictability and parallelism of multitrait adaptation. Journal of Heredity 109:59–70

Langerhans RB & DeWitt TJ (2004) Shared and unique features of evolutionary diversification. American Naturalist 164:335–349

Linn CE, Darnbroski HR, Feder JL, Berlocher SH, Nojima S & Roelofs WL (2004) Postzygotic isolating factor in sympatric speciation in Rhagoletis flies: Reduced response of hybrids to parental host-fruit odors. Proceedings of the National Academy of Sciences of the United States of America 101:17753–17758

Loo WT, Garcia-Loor J, Dudaniec RY, Kleindorfer S & Cavanaugh CM (2019) Host phylogeny, diet, and habitat differentiate the gut microbiomes of Darwin’s finches on Santa Cruz Island. Scientific Reports 9:18781

Martin BT, Douglas MR, Chafin TK, Jr JSP, Birkhead RD, Phillips CA & Douglas ME (2020) Contrasting signatures of introgression in North American box turtle (Terrapene spp.) contact zones. Molecular Ecology 29:4186–4202

Matlab (2017) 9.3.0.713579 (R2017b).

Matsubayashi KW, Ohshima I & Nosil P (2010) Ecological speciation in phytophagous insects. Entomologia Experimentalis Et Applicata 134:1–27

Mendelson TC, Martin MD & Flaxman SM (2014) Mutation-order divergence by sexual selection: Diversification of sexual signals in similar environments as a first step in speciation. Ecology Letters 17:1053–1066

Meyers PJ, Doellman MM, Ragland GJ, Hood GR, Egan SP, Powell THQ, Nosil P & Feder JL (2020) Can the genomics of ecological speciation be predicted across the divergence continuum from host races to species? A case study in Rhagoletis. Philosophical Transactions of the Royal Society of London B-Biological Sciences 375:20190534

Mitter C, Farrell B & Wiegmann B (1988) The phylogenetic study of adaptive zones - Has phytophagy promoted insect diversification? American Naturalist 132:107–128

Nilsson PA, Hulthen K, Chapman B, Hansson LA, Brodersen J, Baktoft H, Vinterstare J, Bronmark C & Skov C (2017) Species integrity enhanced by a predation cost to hybrids in the wild. Biology Letters 13:20170208

Nosil P 2012 Ecological speciation. Oxford University Press, Oxford

Nosil P, Crespi BJ & Sandoval CP (2002) Host-plant adaptation drives the parallel evolution of reproductive isolation. Nature 417:440–443

Nylin S, Slove J & Janz N (2014) Host plant utilization, host range oscillations and diversification in nymphalid butterflies: A phylogenetic investigation. Evolution 68:105–124

Ortega J (in prep) Genomic landscape of Tephritis conura

Pieterse W, Benitez HA & Addison P (2017) The use of geometric morphometric analysis to illustrate the shape change induced by different fruit hosts on the wing shape of Bactrocera dorsalis and Ceratitis capitata (Diptera: Tephritidae). Zoologischer Anzeiger 269:110–116

Porter CK & Benkman CW (2019) Character displacement of a learned behaviour and its implications for ecological speciation. Proceedings of the Royal Society B-Biological Sciences 286:20190761

R Core Team (2019) R: A language and environment for statistical computing.

Rohlf JF (2017) tpsDig2: Digitize coordinates of landmarks and capture outlines.

Rohlf JF (2018) tpsUtil: Utility program useful when working with tps files.

Romstock-Volkl M (1997) Host race formation in Tephritis conura: Determinants from three trophic levels. Ecological Studies 130:21–38

Roriz AKP, Japyassu HF, Caceres C, Vera MT & Joachim-Bravo IS (2019) Pheromone emission patterns and courtship sequences across distinct populations within Anastrepha fraterculus (Diptera-Tephritidae) cryptic species complex. Bulletin of Entomological Research 109:408–417

Rundle HD & Nosil P (2005) Ecological speciation. Ecology Letters 8:336–352

Rundle HD & Rowe L (2018) The contribution of sexual selection to ecological and mutation-order speciation. Evolution 72:2571–2575

Schluter D (1993) Adaptive radiation in sticklebacks - Size, shape and habitat use efficiency. Ecology 74:699–709

Schluter D 2008 The ecology of adaptive radiation. Oxford Univ. Press, Oxford

Schluter D (2009) Evidence for ecological speciation and its alternative. Science 323:737–741

Servedio MR & Noor MAF (2003) The role of reinforcement in speciation: Theory and data. Annual Review of Ecology Evolution and Systematics 34:339–364

Shelomi M (2012) Where are we now? Bergmann’s rule sensu lato in insects. American Naturalist 180:511–519

Sivinski J (2000) Breeding habits and sex in families closely related to the Tephritidae: Opportunities for comparative studies of the evolution of fruit fly behavior. In: Aluja M & Norrbom AL (eds) Fruit Flies (Tephritidae). Boca Raton, pp23–37

Sivinski J & Pereira R (2005) Do wing markings in fruit flies (Diptera: Tephritidae) have sexual significance? Florida Entomologist 88:321–324

Taylor EB, Boughman JW, Groenenboom M, Sniatynski M, Schluter D & Gow JL (2006) Speciation in reverse: morphological and genetic evidence of the collapse of a three-spined stickleback (Gasterosteus aculeatus) species pair. Molecular Ecology 15:343–355

Teets NM, Dias VS, Pierce BK, Schetelig MF, Handler AM & Hahn DA (2019) Overexpression of an antioxidant enzyme improves male mating performance after stress in a lek-mating fruit fly. Proceedings of the Royal Society B-Biological Sciences 286:20190531

Thibert-Plante X & Gavrilets S (2013) Evolution of mate choice and the so-called magic traits in ecological speciation. Ecology Letters 16:1004–1013

Weiner J 1994 The beak of the finch: A story of evolution in our time. Jonathan Cape, London

Wiens JJ, Lapoint RT & Whiteman NK (2015) Herbivory increases diversification across insect clades. Nature Communications 6:8370

Zelditch ML 2004 Geometric morphometrics for biologists. Academic Press, Amsterdam

