## Supplementary material for "Non-parallel morphological divergence following colonization of a new host plant": All supplementary material

Table S1: **Geolocations for sample populations.** CO stands for *Cirsium oleraceum*, CH stands for *Cirsium heterophyllum*, GE stands for Germany, SK represents southern Sweden, ST represents central Sweden, LI stands for Lithuania, ES stands for Estonia and finally FI stands for Finland.

| Population | Name | Latitude | Longitude |
|---|---|---|---|
| COGE | OD7 | 51.112800 | 9.597117 |
| COSK | Trainsite | 55.905000 | 13.413000 |
| CHSK | Ditchsite | 55.864156 | 13.745842 |
| CHST | Grythyttan | 59.628468 | 14.577117 |
| COLI | Strumbres | 55.652206 | 21.938014 |
| COES | Laatre | 57.879930 | 26.248673 |
| CHES | Saaremaa | 58.436099 | 22.937953 |
| CHFI | Nakkila | 61.352485 | 22.056471 |

Supplementary material S1: Fly rearing

Field collection was performed during June and July 2018. All sampling was performed in compliance with the Nagoya protocol. *Tephritis conura* flies were reared in our laboratory at the department of Biology, Lund University. Infested thistles were separated in individual plastic cups, covered with netting. As soon as flies started to emerge, they were given honey diluted in water smeared on top of the netting for sustenance and put on a three-day hold, which was determined to be sufficient time to allow all pupae in the cup to eclose as adults. Subsequently, the cups containing the emerged flies were moved to a climate chamber set to 7 C° and an 8-hour light cycle. A constant supply of honey mixed with water was given to all specimens during containment. In clutches that were large enough we sampled two males and two females and froze them in -80 C°. This allowed us to have a backup fly per sex per family should any of the individuals be damaged or otherwise unusable for further analysis. No more than one fly of each sex per thistle bud was ultimately used in any analysis.

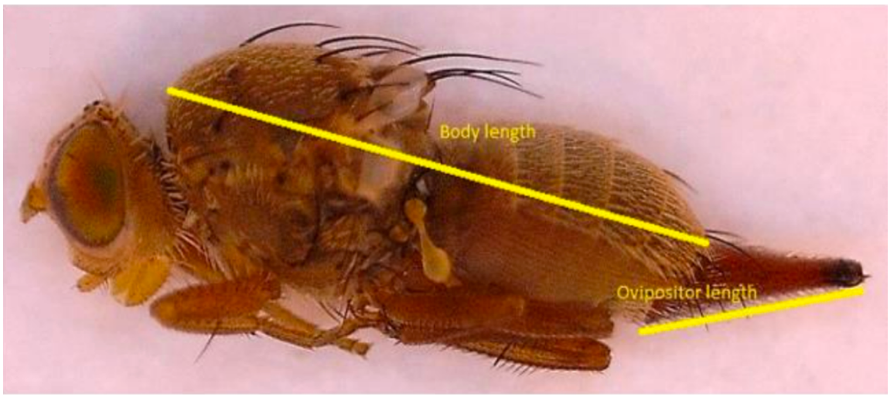


Figure S1: **Body measurements.** All flies measured using the exact same positions.

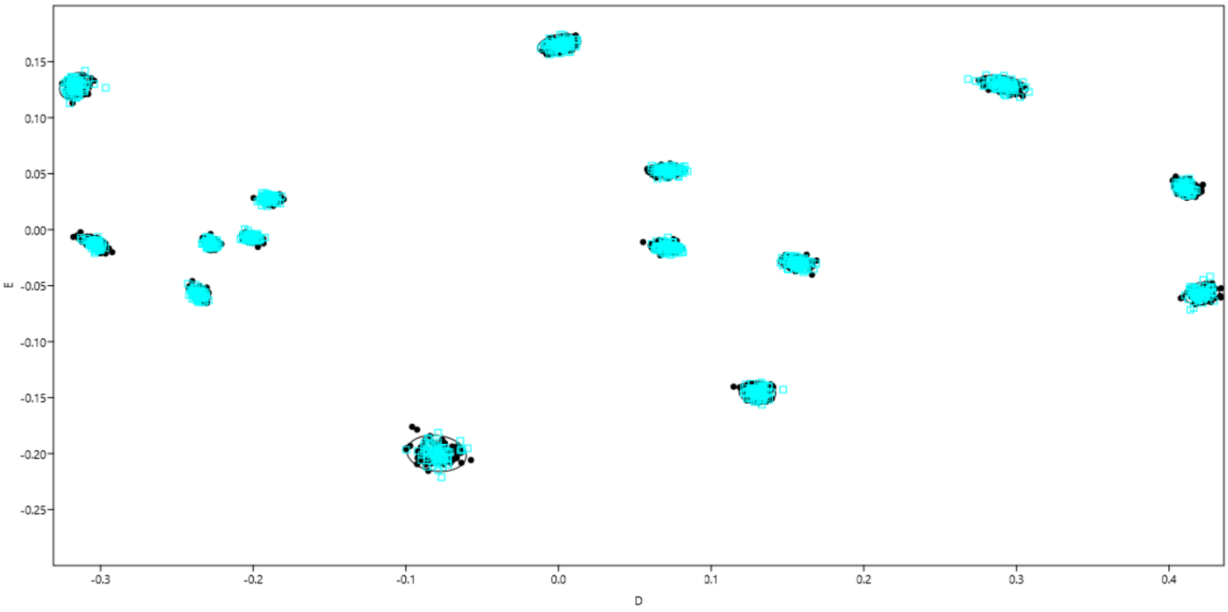


Figure S2: **X and Y coordinates landmarks.** Includes all digitized flies after application of a Procrustes fit.

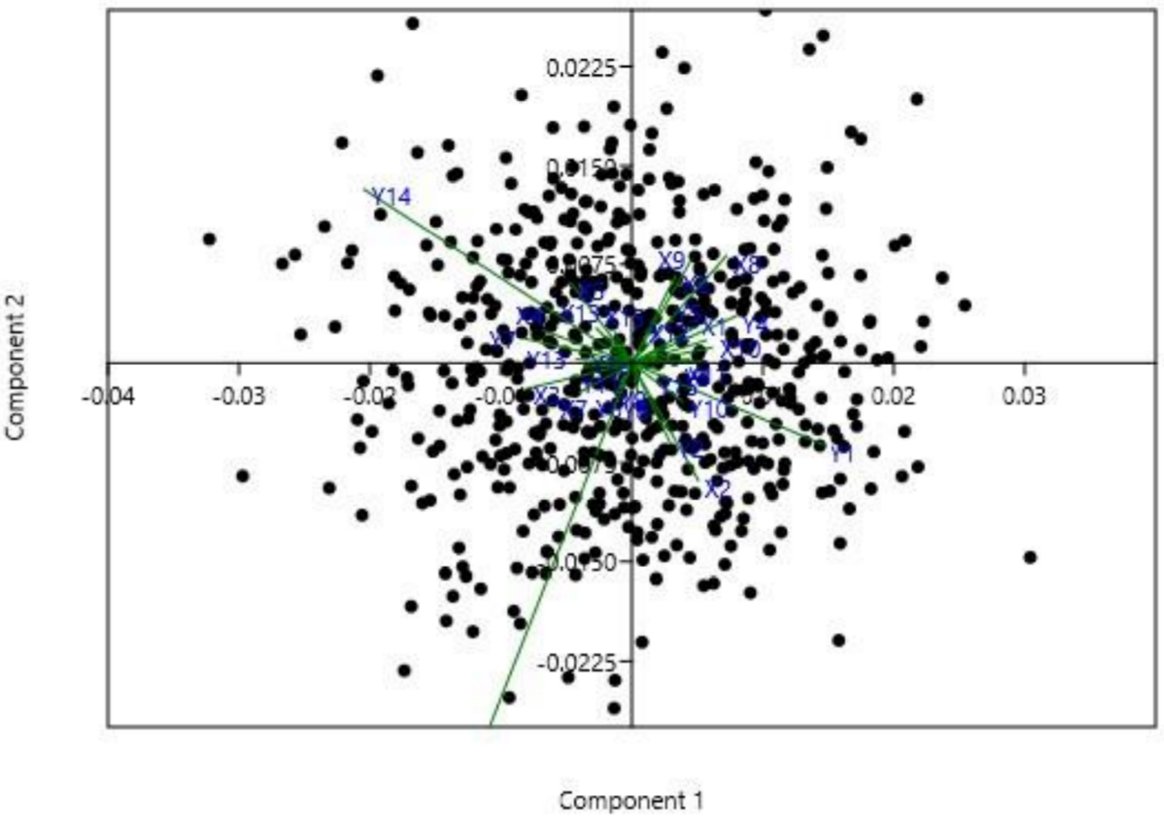


Figure S3: **Principal component analysis of landmark data.** The first two principal components explain 30,89% of all variation. Loadings represents X and Y coordinates of all landmarks.

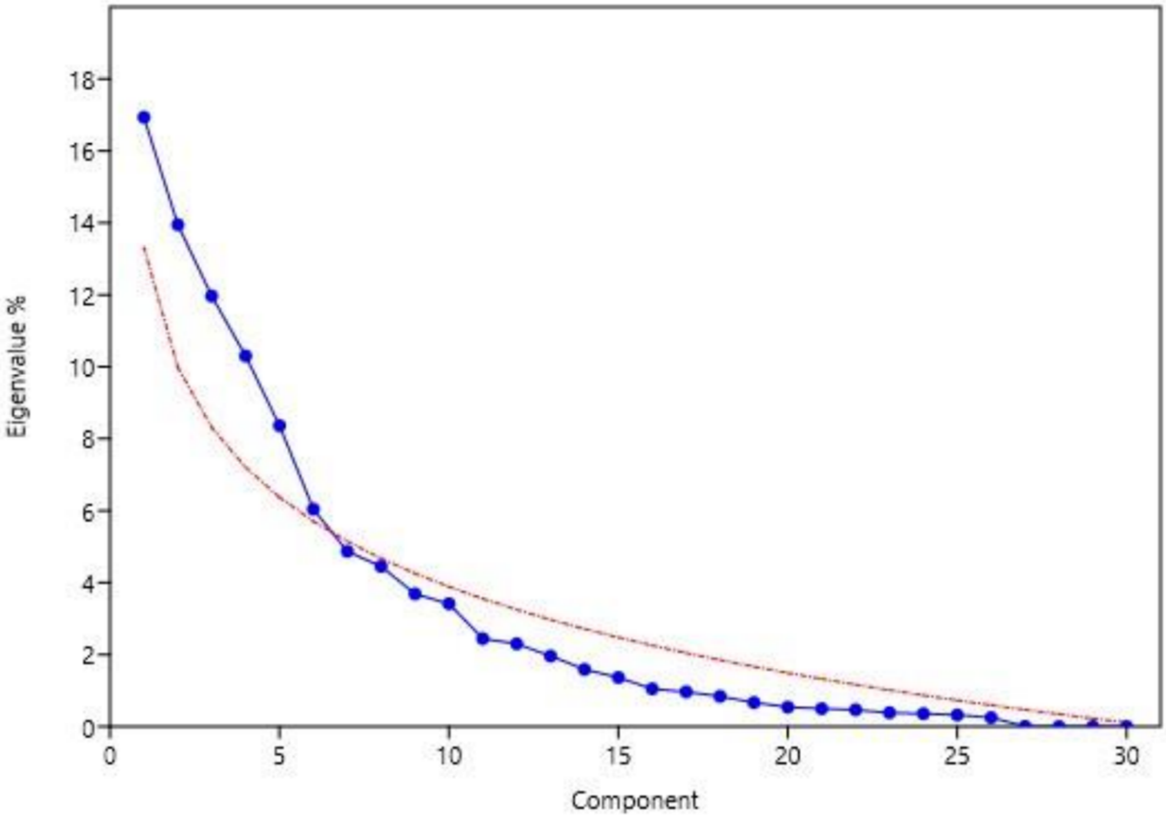


Figure S4: **Scree plot of eigenvalues on landmark data with applied broken stick.** This guided our choice in bringing six principal warps from the landmark analysis to the subsequent MANOVA.

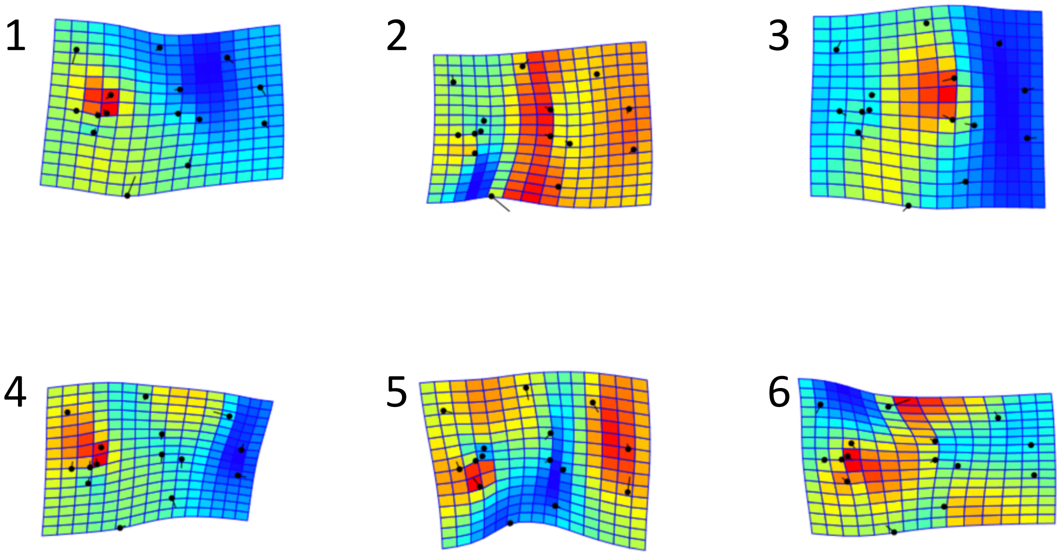


Figure S5: **Visualization of wing shape differentiation in relative warps 1-6.** Heatmap represents degree of local differentiation. The relative warps represent wing shape phenotypes.

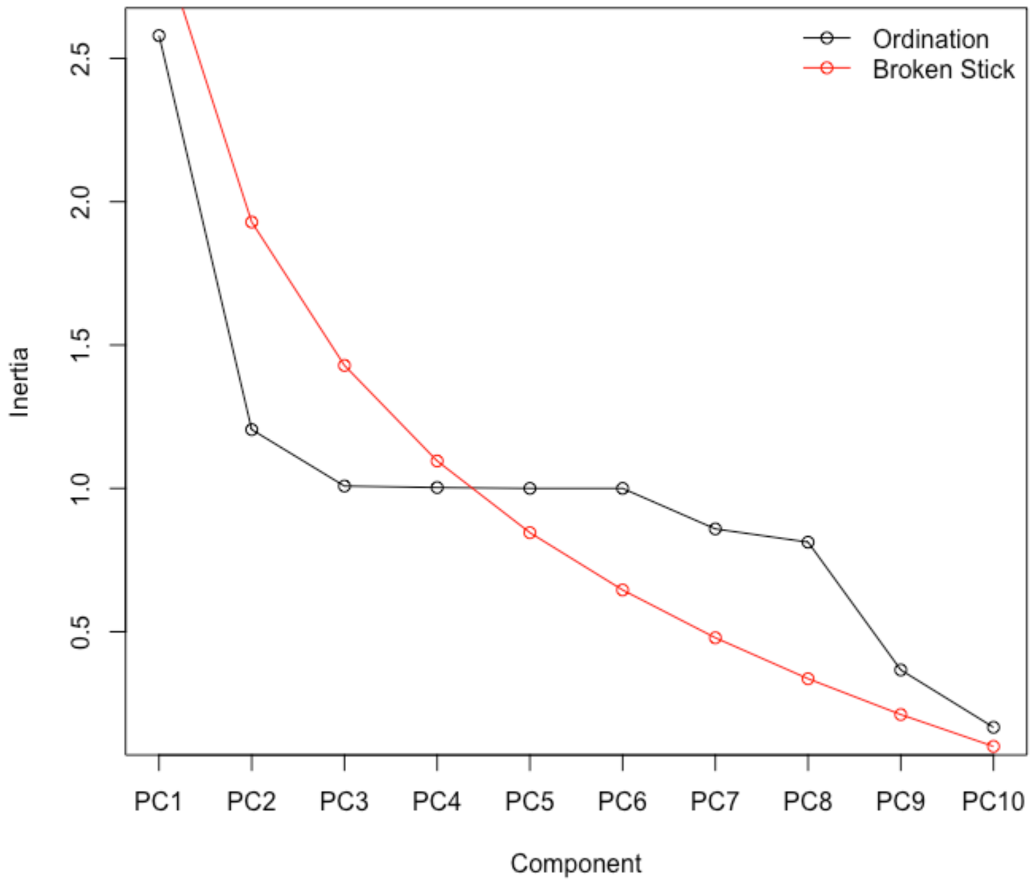


Figure S6: **Scree plot of eigenvalues on fly phenotypic analysis with applied broken stick.** The first four principal components were tested against host race in a general linear model, neither being significant.

Table S2: **Trait loadings on full PCA.** Here including both sexes, hence the exclusion of ovipositor length. LMPC stands for landmark principal component, ie the six wing shape warps.

| | PC1 | PC2 | PC3 | PC4 |
|---|---|---|---|---|
| Body length | 0.518331498 | -0.15741176 | 0.007539775 | 0.05391370 |
| Wing length | 0.557259345 | 0.01131358 | 0.071501627 | -0.03101136 |
| Wing width | 0.559843979 | 0.13817824 | -0.070691887 | -0.02393056 |
| Melanisation ratio | 0.001354099 | 0.73782782 | -0.003092571 | 0.01671527 |
| LMPC1 | 0.070151748 | -0.37286993 | -0.565677571 | 0.10449625 |
| LMPC2 | -0.068587949 | 0.14092765 | -0.123522891 | -0.54689363 |
| LMPC3 | 0.234588128 | -0.06784420 | 0.590428583 | -0.01762155 |
| LMPC4 | 0.166935931 | -0.05976643 | -0.435516436 | -0.24360949 |
| LMPC5 | -0.120792296 | -0.46251963 | 0.299504584 | 0.08124771 |
| LMPC6 | -0.015868792 | -0.17498159 | 0.163068300 | -0.78677197 |

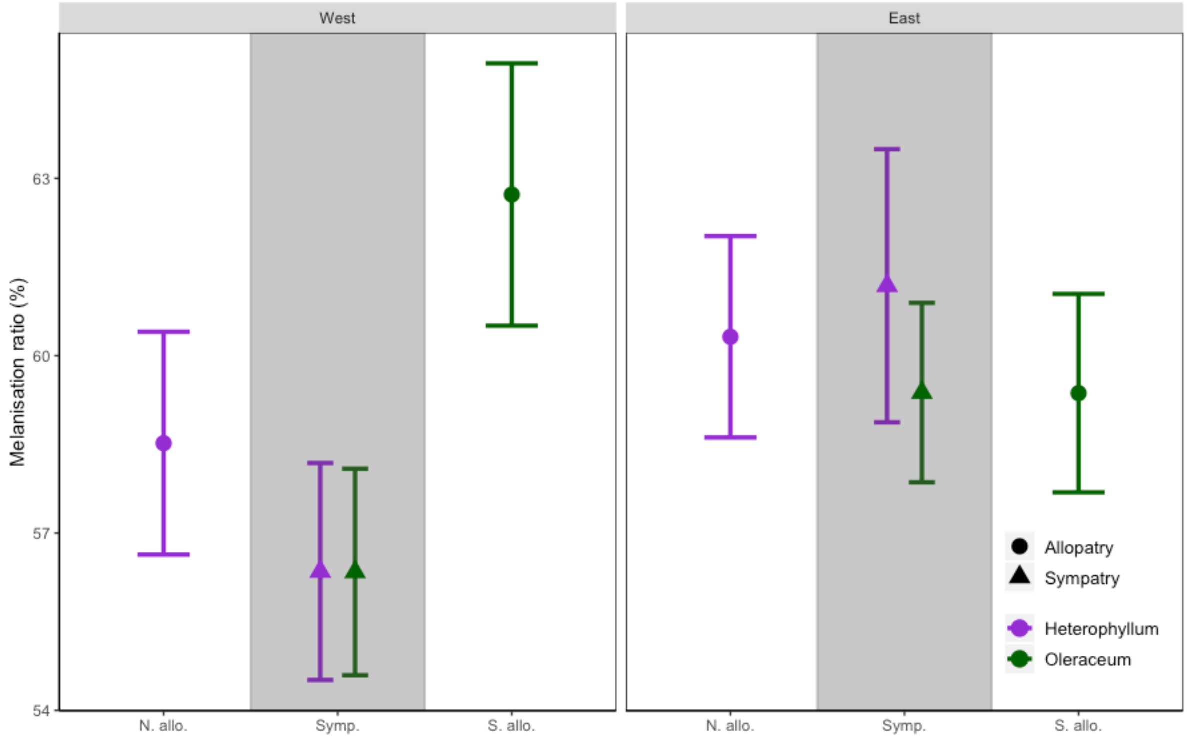


Figure S7: **Mean trait values of melanisation ratio with 95% confidence intervals of the mean.** Only female *T. conura* are included. Mean values of *T. conura* melanisation ratio separated by population. 'West' and 'East' represents from which side of the Baltic Sea the population is sampled. 'N. allo.' stands for Northern allopatry, 'Symp.' stands for sympatry and 'S. allo.' stands for Southern allopatry.

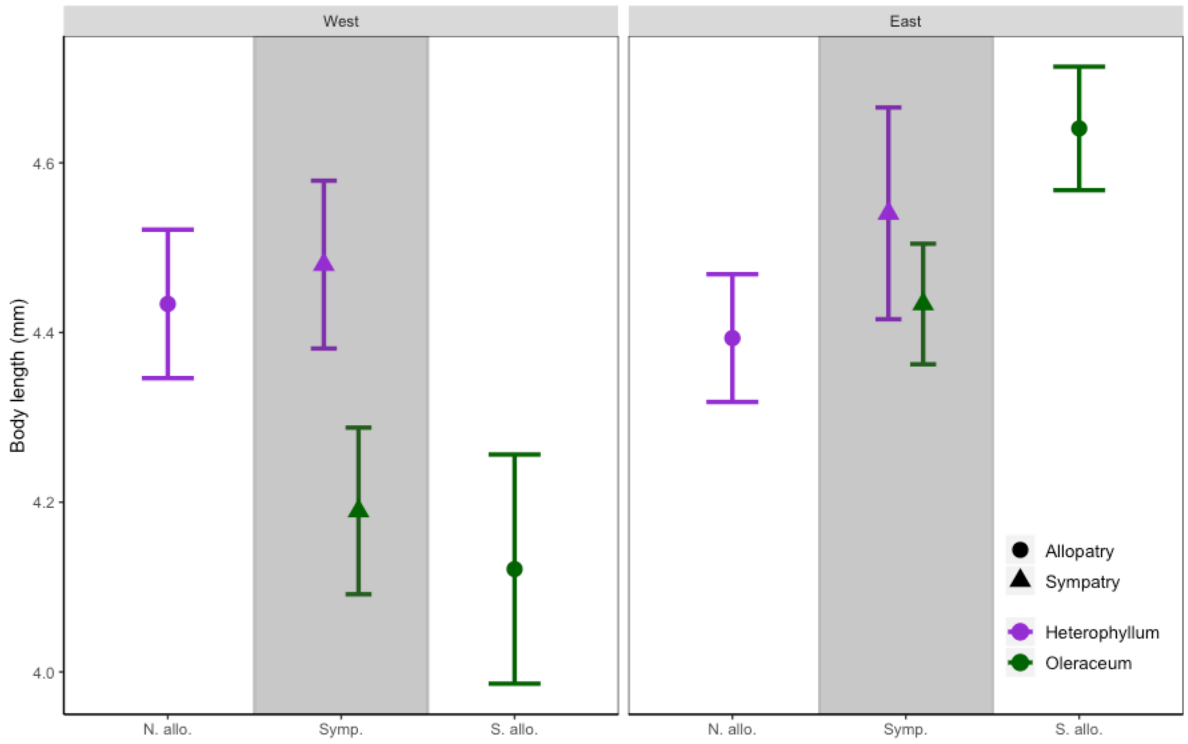


Figure S8: **Mean trait values of body length with 95% confidence intervals of the mean.** Here only male *T. conura* are included. Mean values of *T. conura* body length separated by population. 'West' and 'East' represents from which side of the Baltic Sea the population is sampled. 'N. allo.' stands for Northern allopatry, 'Symp.' stands for sympatry and 'S. allo.' stands for Southern allopatry.

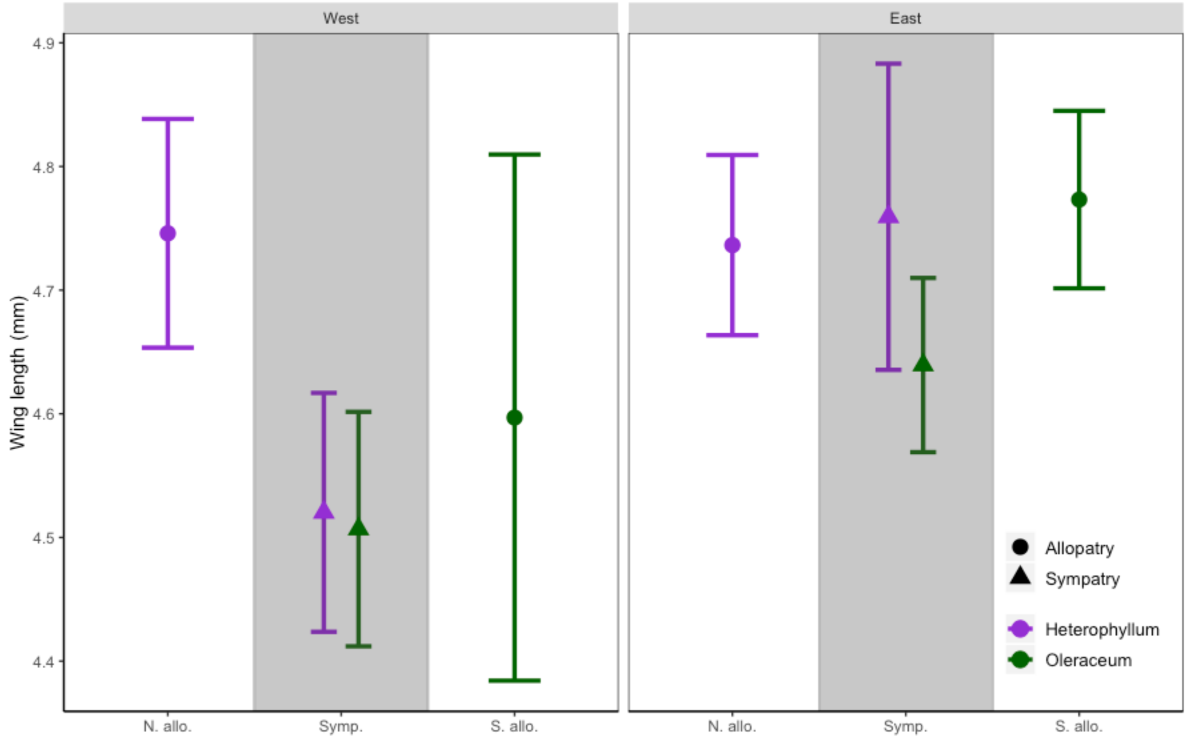


Figure S9: **Mean trait values of wing length with 95% confidence intervals**: Only male *T. conura* are included. Mean values of *T. conura* wing length separated by population. 'West' and 'East' represents from which side of the Baltic Sea the population is sampled. 'N. allo.' stands for Northern allopatry, 'Symp.' stands for sympatry and 'S. allo.' stands for Southern allopatry.

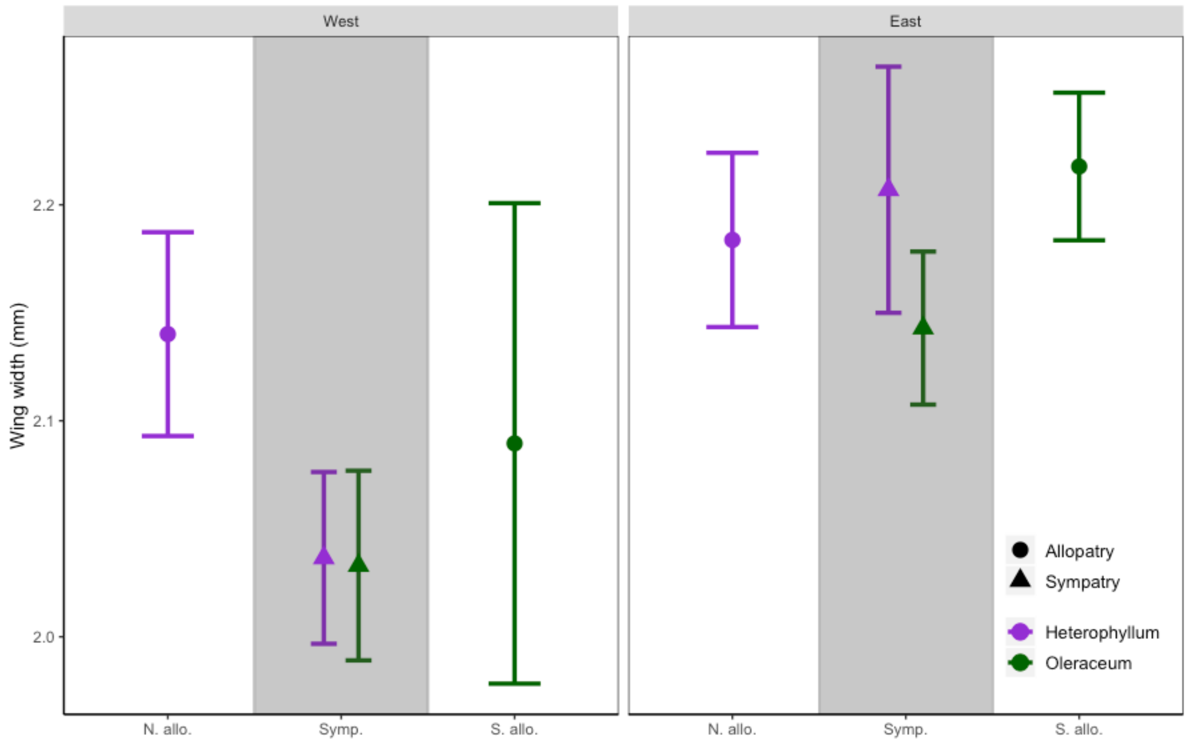


Figure S10: **Mean trait values of wing width with 95% confidence intervals**: Only male *T. conura* are included. Mean values of *T. conura* wing length separated by population. 'West' and 'East' represents from which side of the Baltic Sea the population is sampled. 'N. allo.' stands for Northern allopatry, 'Symp.' stands for sympatry and 'S. allo.' stands for Southern allopatry.

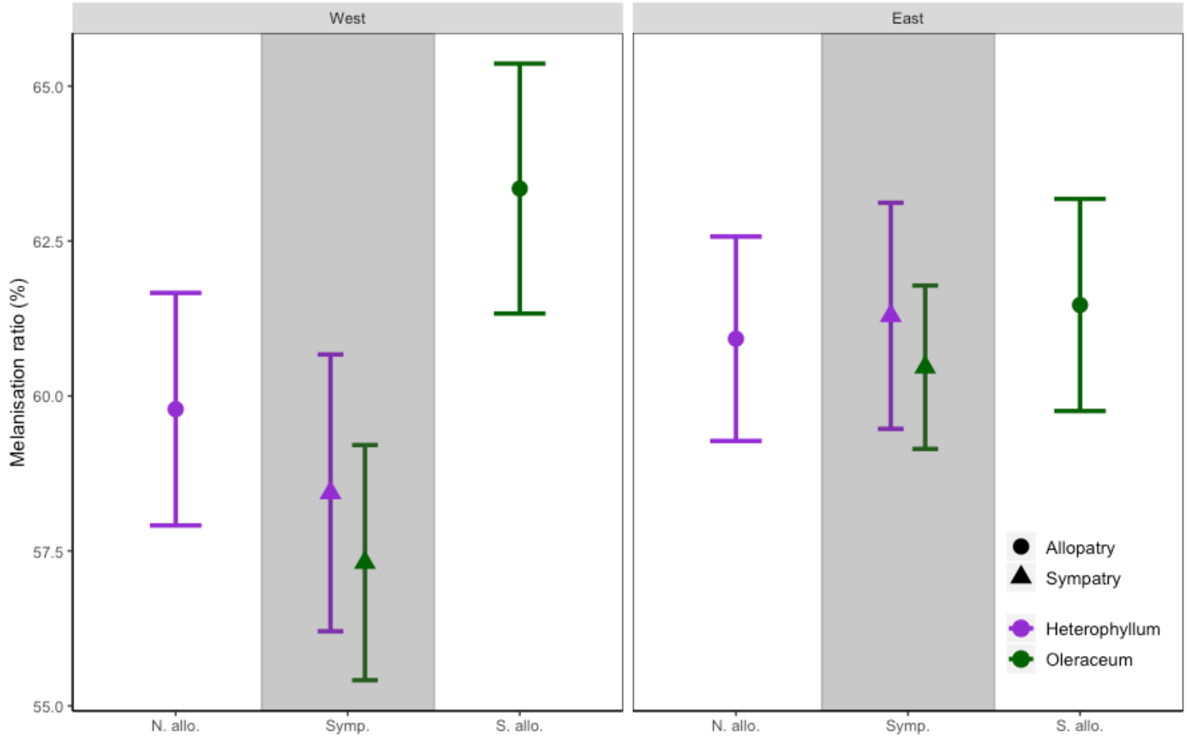


Figure S11: **Mean trait values of wing melanisation with 95% confidence intervals**: Only male *T. conura* are included. Mean values of *T. conura* wing length separated by population. ‘West’ and ‘East’ represents from which side of the Baltic Sea the population is sampled. ‘N. allo.’ stands for Northern allopatry, ‘Symp.’ stands for sympatry and ‘S. allo.’ stands for Southern allopatry.

Table S3: **Post-Hoc ANOVA.** Only traits that differed significantly between groups are included.

| Independent | Dependent | F | num df | den df | P | |
|---|---|---|---|---|---|---|
| Females | | | | | | |
| Host plant | Body length | 4.29 | 1 | 277 | 0.039 | * |
| | Ovipositor length | 50.65 | 1 | 277 | < 0.001 | *** |
| | Wing length | 4.08 | 1 | 277 | 0.044 | * |
| | Wing shape, warp 3 | 4.43 | 1 | 277 | 0.036 | * |
| | Wing shape, warp 5 | 10.63 | 1 | 277 | 0.001 | ** |
| Co-existence | Ovipositor length | 14.39 | 1 | 277 | < 0.001 | *** |
| | Wing length | 9.92 | 1 | 277 | 0.002 | ** |
| | Wing width | 20.61 | 1 | 277 | < 0.001 | *** |
| | Melanisation ratio | 9.83 | 1 | 277 | 0.002 | ** |
| | Wing shape, warp 2 | 10.98 | 1 | 277 | 0.001 | ** |
| | Wing shape, warp 3 | 7.62 | 1 | 277 | 0.006 | ** |
| | Wing shape, warp 5 | 14.45 | 1 | 277 | < 0.001 | *** |
| Baltic | Body length | 24.96 | 1 | 277 | < 0.001 | *** |
| | Ovipositor length | 28.23 | 1 | 277 | < 0.001 | *** |
| | Wing length | 24.73 | 1 | 277 | < 0.001 | *** |
| | Wing width | 44.34 | 1 | 277 | < 0.001 | *** |
| | Melanisation ratio | 6.02 | 1 | 277 | 0.015 | * |
| | Wing shape, warp 1 | 5.14 | 1 | 277 | 0.024 | * |
| | Wing shape, warp 3 | 26.32 | 1 | 277 | < 0.001 | *** |
| | Wing shape, warp 4 | 18.52 | 1 | 277 | < 0.001 | *** |
| | Wing shape, warp 5 | 33.41 | 1 | 277 | < 0.001 | *** |
| Host plant x co-existence | Body length | 10.68 | 1 | 277 | 0.001 | ** |
| | Wing width | 10.5 | 1 | 277 | 0.001 | ** |
| | Wing shape, warp 1 | 8.97 | 1 | 277 | 0.003 | ** |
| | Wing shape, warp 3 | 11.16 | 1 | 277 | < 0.001 | *** |

| | | | | | | |
|---|---|---|---|---|---|---|
| | Wing shape, warp 5 | 11.97 | 1 | 277 | < 0.001 | *** |
| Host plant x Baltic | Body length | 24.24 | 1 | 277 | < 0.001 | *** |
| | Ovipositor length | 15.84 | 1 | 277 | < 0.001 | *** |
| | Wing shape, warp 3 | 15.61 | 1 | 277 | < 0.001 | *** |
| Co-existence x Baltic | Ovipositor length | 5.44 | 1 | 277 | 0.02 | * |
| | Melanisation ratio | 11.05 | 1 | 277 | 0.001 | ** |
| | Wing shape, warp 5 | 19.96 | 1 | 277 | < 0.001 | *** |
| Three-way interaction | Body length | 7.40 | 1 | 277 | 0.007 | ** |
| | Ovipositor length | 4.33 | 1 | 277 | 0.038 | * |
| | Wing width | 5.02 | 1 | 277 | 0.026 | * |
| | Wing shape, warp 3 | 5.57 | 1 | 277 | 0.019 | * |
| | Wing shape, warp 4 | 12.94 | 1 | 277 | < 0.001 | *** |
| Males | | | | | | |
| Host plant | Wing shape, warp 5 | 8.61 | 1 | 280 | 0.004 | ** |
| Co-existence | Wing length | 14.55 | 1 | 280 | < 0.001 | *** |
| | Wing width | 16.52 | 1 | 280 | < 0.001 | *** |
| | Melanisation ratio | 7.37 | 1 | 280 | 0.007 | ** |
| | Wing shape, warp 2 | 12.75 | 1 | 280 | < 0.001 | *** |
| | Wing shape, warp 3 | 9.52 | 1 | 280 | 0.002 | ** |
| | Wing shape, warp 5 | 7.42 | 1 | 280 | 0.007 | ** |
| Baltic | Body length | 24.98 | 1 | 280 | < 0.001 | *** |
| | Wing length | 11.51 | 1 | 280 | < 0.001 | *** |
| | Wing width | 38.74 | 1 | 280 | < 0.001 | *** |
| | Melanisation ratio | 5.69 | 1 | 280 | 0.018 | * |
| | Wing shape, warp 1 | 5.11 | 1 | 280 | 0.025 | * |
| | Wing shape, warp 3 | 15.48 | 1 | 280 | < 0.001 | *** |
| | Wing shape, warp 4 | 15.78 | 1 | 280 | < 0.001 | *** |
| | Wing shape, warp 5 | 50.31 | 1 | 280 | < 0.001 | *** |
| | Wing shape, warp 6 | 0.07 | 1 | 280 | 0.003 | ** |

| | | | | | | |
|---|---|---|---|---|---|---|
| Host plant x Co-existence | Body length | 13.62 | 1 | 280 | < 0.001 | *** |
| | Wing shape, warp 1 | 5.46 | 1 | 280 | 0.02 | * |
| | Wing shape, warp 3 | 9.83 | 1 | 280 | 0.002 | ** |
| | Wing shape, warp 6 | 7.39 | 1 | 280 | 0.007 | ** |
| Host plant x Baltic | Body length | 29.47 | 1 | 280 | 29.47 | |
| | Wing shape, warp 3 | 3.95 | 1 | 280 | 0.045 | * |
| | Wing shape, warp 5 | 4.62 | 1 | 280 | 0.033 | * |
| Co-existence x Baltic | Melanisation ratio | 5.85 | 1 | 280 | 0.016 | * |
| | Wing shape, warp3 | 5.42 | 1 | 280 | 0.021 | * |
| | Wing shape, warp 5 | 8.34 | 1 | 280 | 0.004 | ** |
| Three-way interaction | Body length | 7.96 | 1 | 280 | 0.005 | ** |
| | Wing length | 4.47 | 1 | 280 | 0.035 | * |
| | Wing width | 4.49 | 1 | 280 | 0.035 | * |
| | Wing shape, warp 3 | 4.41 | 1 | 280 | 0.037 | * |
| | Wing shape, warp 5 | 9.49 | 1 | 280 | 0.002 | ** |

Table S4: **Nested MANOVA results**.

| Test for | Factor | F | df | P | Partial variance explained (%) |
|---|---|---|---|---|---|
| Shared divergence | Host Plant | 4.96 | 11.273 | <0.001 | 30.6 |
| Unique histories | Transect | 6.85 | 11.273 | <0.001 | 37.8 |
| Unique divergence | Population (Transect x Host plant) | 3.81 | 11.273 | <0.001 | 25.3 |

Table S5: **Bootstrap loadings LDA**. Cell color represents whether bootstrapped estimate of each trait is greater or smaller than standard error. Bootstrap performed with 100 000 replications.

| | Discriminant axis 1, East | Discriminant axis 1, West | Discriminant axis 2, East | Discriminant axis 2, West |
|---|---|---|---|---|
| Body Length | 3,87 ± 0,94 | -0,84 ± 2,03 | -2,02 ± 1,13 | -2,97 ± 1,12 |
| Ovipositor Length | -4,4 ± 1,33 | -4,95 ± 3,23 | -0,02 ± 1,64 | 5,73 ± 3,83 |
| Wing Length | -2,21 ± 0,93 | 0,04 ± 1,09 | 1,6 ± 1,18 | 1,1 ± 0,95 |
| Wing Width | -2,65 ± 1,91 | 3,29 ± 2,45 | -1,38 ± 2,38 | 1,29 ± 2,3 |
| Melanisation Ratio | 0,03 ± 0,03 | 0,03 ± 0,04 | -0,06 ± 0,06 | 0,02 ± 0,03 |
| PC1 | -25,40 ± 19,39 | 13,69 ± 19,34 | -44,68 ± 19,85 | -7,26 ± 20,14 |
| PC2 | 12,27 ± 18,33 | 14,81 ± 22,29 | -41,04 ± 22,2 | -19,02 ± 23,5 |
| PC3 | -14,93 ± 21,80 | -54,47 ± 30,69 | -44,45 ± 30,64 | -14,56 ± 32,83 |
| PC4 | 14,74 ± 32,09 | -40,87 ± 27,05 | -70,63 ± 26,89 | -10,34 ± 30,8 |
| PC5 | 133,06 ± 22,72 | 54,53 ± 41,3 | 12,02 ± 36,76 | 30,83 ± 33,99 |
| PC6 | 39,17 ± 27,2 | 8,84 ± 27,72 | -6,02 ± 37,15 | 3,3 ± 29,78 |
